# Biogeographic study of human gut-associated crAssphage suggests impacts from industrialization and recent expansion

**DOI:** 10.1101/384677

**Authors:** Tanvi P. Honap, Krithivasan Sankaranarayanan, Stephanie L. Schnorr, Andrew T. Ozga, Christina Warinner, Cecil M. Lewis

## Abstract

CrAssphage (cross-assembly phage) is a bacteriophage that was first discovered in human gut metagenomic data. CrAssphage belongs to a diverse family of crAss-like bacteriophages thought to infect gut commensal bacteria belonging to *Bacteroides* species. However, not much is known about the biogeography of crAssphage and whether certain strains are associated with specific human populations. In this study, we screened publicly available human gut metagenomic data from 3,341 samples for the presence of crAssphage *sensu stricto* (NC_024711.1). We found that crAssphage prevalence is low in traditional, hunter-gatherer populations, such as the Hadza from Tanzania and Matses from Peru, as compared to industrialized, urban populations. Statistical comparisons showed no association of crAssphage prevalence with variables such as age, sex, body mass index, and health status of individuals. Phylogenetic analyses show that crAssphage strains reconstructed from the same individual over multiple time-points, cluster together. CrAssphage strains from individuals from the same study population do not always cluster together. Some evidence of clustering is seen at the level of broadly defined geographic regions, however, the relative positions of these clusters within the crAssphage phylogeny are not well-supported. We hypothesize that this lack of strong biogeographic structuring is suggestive of a recent expansion event within crAssphage. Using a Bayesian dating approach, we estimate this expansion has occurred within the past 200 years. Overall, we determine that crAssphage presence is associated with an industrialized lifestyle. The absence of strong biogeographic structuring within global crAssphage strains is likely due to a recent population expansion within this bacteriophage.

## Introduction

The virome harbors the most abundant and diverse set of genes on earth, with deep impacts on host biology, including the very architecture of host genomes [1]. Often described as the “dark matter” of biology [2], the virome has an underexplored role in human health [3].

Much of the human gut virome is composed of double-stranded DNA bacteriophages [4], which are argued to be the primary regulators of bacterial biomass [5]. Bacteriophages have known impacts on the human gut; for example, they are involved in the transmission of bacterial antibiotic resistance genes, and facilitate bacterial carbohydrate utilization [4]. The majority of the bacteriophages found in the human gut belong to the families *Siphoviridae, Podoviridae,* and *Myoviridae* within the order Caudovirales [6]; however, our knowledge of the true bacteriophage diversity is limited [7]. While the human virome is a component of the human microbiome, it is not as amendable to high throughput taxonomic characterization as the archaeal and bacterial subsets. Specifically, viruses lack “universal” taxonomically diagnostic marker genes (for example, the 16S ribosomal RNA gene in archaea and bacteria), preventing characterization using current amplicon-based sequencing strategies.

Recent advances in metagenomic sequencing and development of data mining tools now allow for large-scale studies of human gut bacteriophage diversity using shotgun-sequencing data from human fecal samples [7]. From such data, one particular DNA bacteriophage, crAssphage, appears to be remarkably abundant in the human gut [8]. CrAssphage (cross-assembly phage) was named after the crAss (cross-assembly) software originally used to discover the bacteriophage [8]. CrAssphage is prevalent in available gut metagenomic data from the U.S. [9], comprising up to 90% of viral particle-derived metagenomes and up to 22% of reads in a total fecal community from the U.S. National Institutes of Health’s Human Microbiome Project cohort [10]. Moreover, crAssphage is up to six times more abundant than all other known bacteriophages that could be reconstructed from these publicly available metagenomes [8]. Bacteria belonging to phylum Bacteroidetes, which are commonly found gut commensals, were predicted to be the host of this bacteriophage [8].

A recent study has shown that crAssphage, which is now referred to as crAssphage *sensu stricto,* is actually a member of a diverse group of crAss-like bacteriophages [11]. The first isolated member of this group, ΦCrAss001, shows podovirus-like morphology and infects the bacterium, *Bacteroides intestinalis* [12]. In addition to the human gut, crAss-like phages are found in nonhuman primate and termite guts, terrestrial and groundwater sources, and oceanic environments [11, 13–15]. Across metagenomic studies, crAssphage *sensu stricto* is nearly exclusively associated with the human gut microbiome [8, 11] and has been proposed as a potential biomarker for fecal contamination [16–20].

Recent research suggests that crAssphage may be present in infants as early as one-month after birth [21], which is partly explained by the fact that *Bacteroides* species are among the most abundant members of the gut microbiomes of newborns [22]. It has also been suggested that while most adults contain a single dominant crAssphage strain [21], a minority can carry multiple (hundreds) crAssphage strains [20]. However, the means by which crAssphage is initially acquired [13, 21], strain variation within an individual [20], overall geographic distribution [20, 23], and prevalence in other environmental reservoirs has not been fully elucidated. Although crAssphage has been detected globally using PCR-based assays [13, 20], a systematic analysis of the strain biogeography of crAssphage is necessary, considering the diverse range of currently reported crAssphage strains and the existence of other crAss-like phages in the human gut [11] that may confound crAssphage detection using PCR-based assays.

Here, we report strain-level prevalence and diversity of crAssphage *sensu stricto* observed across 3,341 gut metagenomic samples originating from globally distributed human populations (Table 1). In this study, we consider genetically non-identical crAssphage sequences as different crAssphage strains [24]. Further, we evaluate crAssphage prevalence, abundance, and strain diversity as a function of biogeography, human dietary lifestyle and variables including age, sex, body mass index (BMI), and health status.

**Table 1.** Publicly available gut metagenomic datasets used in this study

| Dataset | Description | Samples | Individuals | Accession | Reference |
| --- | --- | --- | --- | --- | --- |
| BCK | Urban mother-infant pairs from Sweden | 400 | 200 | ERP005989 | [22] |
| CNA | Urban individuals of Cheyenne, Arapaho, and non-native ancestry from Oklahoma, USA | 61 | 61 | PRJNA299502 | [25] |
| HAD | Hadza hunter-gatherers from Tanzania | 67 | 67 | SRP056480, SRP110665 | [26], [27] |
| HMP | Urban individuals from USA | 204 | 123 | phs000228.v3.p1 | [10] |
| ITA | Urban individuals from Italy | 11 | 11 | SRP056480 | [26] |
| ISR | Urban individuals from Israel | 950 | 851 | PRJEB11532 | [28] |
| KRL | Urban individuals from Sweden | 145 | 145 | ERP002469 | [29] |
| LIU | Traditional pastoralists from Mongolia | 110 | 110 | SRP080787 | [30] |
| MAT | Matses hunter-gatherers from Peru | 25 | 25 | PRJNA268964 | [31] |
| MHC | Urban individuals from China | 363 | 363 | SRA045646, SRA050230 | [32] |
| MHE | Urban individuals from Sweden and Denmark | 756 | 606 | ERP003612, ERP004605, ERP002061 | [33], [34], [35] |
| XIE | Urban twin-pairs from the UK | 249 | 249 | ERP010708 | [36] |

## Results and discussion

### CrAssphage screening

Publicly available gut metagenome data from 3,341 samples were screened for the presence of crAssphage *sensu stricto* using a two-pronged approach: first, a direct reference-based mapping to crAssphage (NC_024711.1) using Bowtie2 [37], and second, a *de novo* assembly using MEGAHIT [38] followed by a protein-protein BLAST search (BLASTP) [39] against the crAssphage proteins (see Methods). Mapping and assembly statistics are provided in S1 Table.

Using the reference-based mapping approach, we identified 614 samples wherein a high-quality crAssphage strain could be reconstructed (>70% of the genome at >10-fold mean coverage). However, this approach is suitable only for samples containing a single dominant crAssphage strain. 60 samples showed more than 100 heterozygous sites (which corresponds to 0. 1% of the total genome), with the maximum being 1,118 heterozygous sites in a single sample. In order to determine whether the samples comprised multiple crAssphage strains, we performed a *de novo* assembly of each metagenome. We used BLASTP to query the predicted open reading frames (ORFs) for each sample against the crAssphage reference (NC_024711.1) proteins. The BLASTP hits were then filtered to include only those which showed 95% query coverage and 95% identity to crAssphage proteins. These criteria were chosen to avoid false positive hits from crAss-like phages, since the average pairwise identity between members of the crAss-like phage family ranges from 20-40% [11, 40]. Using this approach, we identified 963 samples wherein at least one crAssphage protein was recovered (S2 Table). We were not able to recover all 90 proteins from a single sample, with the maximum number of crAssphage proteins recovered from a single sample being 67. We hypothesize that this is due to the limitations of our *de novo* metagenomic assembly as well as the stringency of our BLASTP search parameters, wherein our approach is likely biased towards avoiding false positive hits. However, our approach is not biased towards recovery of specific crAssphage proteins, since all 90 crAssphage reference proteins were recovered across all samples. Since we were only able to recover a maximum of 67 proteins for a single sample, we decided to use a 50% coverage threshold to call a sample “crAss-positive”, i.e. at least 33 crAssphage proteins had to be recovered from the sample. Using this criterion, 719 samples were found to be crAss-positive. Further, we determined that 497 samples had at least one crAssphage reference protein matched by multiple sample ORFs, suggesting presence of multiple strains.

### crAssphage prevalence

To assess the prevalence of crAssphage among individuals from each study population as well as potential associations with health, individuals were divided into categories based on health status (Table 2). For some datasets (BCK, HMP, ISR, and MHE), multiple samples were available from the same individual; in this case, an individual was considered crAss-positive if at least one sample from that individual was positive. None of the HAD and ITA samples, acquired from Hadza individuals from Tanzania (N=67) and urban Italians (N=11), respectively, were crAss-positive. For the MAT samples acquired from Matses individuals from Peru (N=25), only two were crAss-positive. The Hadza peoples from Tanzania as well as the Matses from the Amazon jungle of Peru have a subsistence practice primarily based on hunting and gathering [31, 41]. Among healthy, urban individuals leading an industrialized lifestyle, crAssphage prevalence ranged from 14.0 % among Chinese individuals from the MetaHIT cohort (MHC; N=185) to 35.7 % among U.S. residents from the HMP cohort (N=123). CrAssphage prevalence showed no significant associations with human health status (Table 2).

**Table 2.**
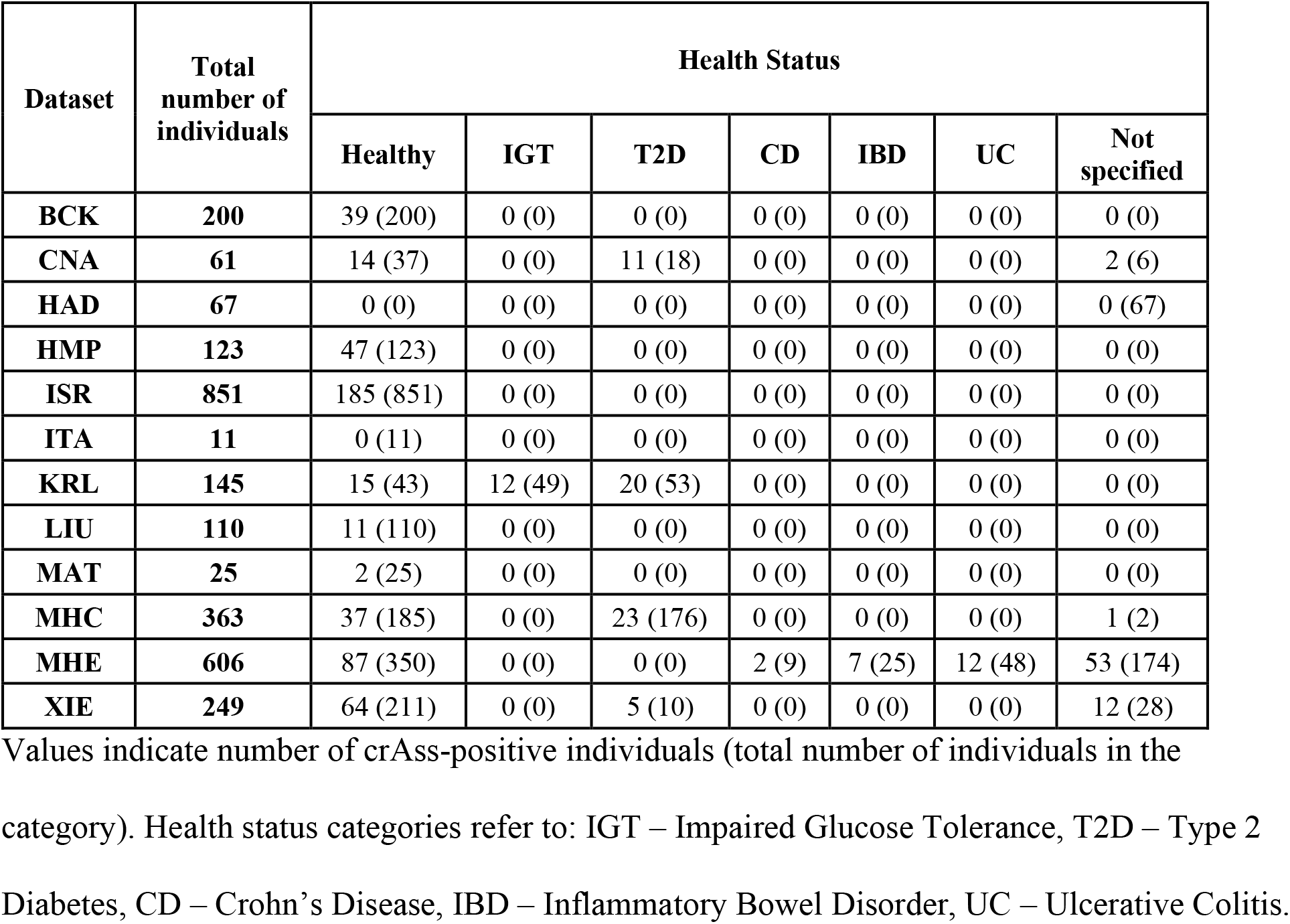
Association of health status with prevalence of crAssphage

Our findings suggest that crAssphage prevalence may be associated with an industrialized lifestyle. A similar finding was reported in a manuscript that was published while our manuscript was under review [15]. This latter study found that gut metagenomic data for rural Malawi and Amazonas individuals from Venezuela [42] as well as the ~5300-year old Tyrolean Iceman [43] show very low numbers of crAssphage sequences. The association of crAssphage with industrialized practices can also be partially explained by its putative bacterial host, *Bacteroides* species [8]. CrAss-like phages can infect *Bacteroides* [44], although it is not known if they are limited to this host genus [45]. Gut microbiome studies have shown that *Bacteroides* tend to be more abundant in industrialized human populations as compared to traditional peoples [46], and the Hadza and Matses individuals are consistent with this pattern [31, 41]. However, the two crAss-positive Matses individuals do not show higher relative abundance of *Bacteroides* as compared to crAss-negative Matses individuals (S1 Fig).

For healthy individuals, we also analyzed metadata, such as age, sex, and BMI, to study potential associations with prevalence of crAssphage. The HAD, ISR, ITA, LIU, and MAT datasets were excluded from this analysis due to unavailability of metadata or insufficient number of crAss-positive individuals. There were no statistically significant differences in the prevalence of crAssphage among individuals based on age, sex, or BMI (Table 3).

**Table 3.** Association of age, sex, and BMI of individuals with prevalence of crAssphage

| Dataset | CNA | HMP | KRL | MHC | MHE | XIE |
| --- | --- | --- | --- | --- | --- | --- |
| **Age** |  |  |  |  |  |  |
| **18 - 40** | 9 (23) | 47 (123) | 0 (0) | 16 (74) | 11 (35) | 1 (4) |
| **41 - 65** | 4 (12) | 0 (0) | 0 (0) | 19 (103) | 47 (215) | 44 (134) |
| **> 65** | 1 (2) | 0 (0) | 15 (43) | 2 (7) | 2 (19) | 19 (73) |
| **Sex** |  |  |  |  |  |  |
| **Male** | 5 (18) | 30 (65) | 0 (0) | 19 (95) | 31 (128) | 0 (0) |
| **Female** | 9 (19) | 17 (58) | 15 (43) | 18 (90) | 29 (142) | 64 (211) |
| **BMI** |  |  |  |  |  |  |
| **< 18.5** | 0 (0) | 0 (0) | 1 (1) | 2 (17) | 0 (3) | 0 (4) |
| **18.5-24.9** | 2 (13) | 29 (68) | 6 (18) | 17 (104) | 29 (130) | 35 (108) |
| **>= 25.0** | 12 (24) | 18 (55) | 8 (24) | 18 (64) | 58 (207) | 29 (98) |

Values denote number of crAss-positive individuals (total number of individuals in the category)

### Phylogenetic analyses

To study the phylogenetic relationships between the crAssphage strains, we selected ten genes which are present in all members of the crAss-like phage family [11]. These genes include those encoding the five putative capsid proteins, a single-stranded DNA-binding protein, a DNA-G family primase, a PD-(D/E)XK family nuclease, and two hypothetical proteins. We then identified a subset of 232 samples wherein only one sample ORF matched these crAssphage reference proteins, suggesting the presence of a single strain of crAssphage in the samples. Furthermore, we confirmed that these ORFs had similar depths of coverage to verify that they represent the same viral genome. We also included the reference crAssphage genome (NC_024711.1), resulting in a total of 233 taxa. A nucleotide alignment comprising these ten genes contained 12,642 sites. A Maximum Likelihood (ML) tree of the 233 strains based on this multi-gene alignment is given in S2 Fig.

According to the ML tree, crAssphage strains from the same individual cluster together. While strains from individuals belonging to the same population/study dataset do cluster together, many such crAssphage clusters are found in each population. Unfortunately, the relative positions of these clusters could not be well-established due to insufficient bootstrap support. Interestingly, we found that the strain from a crAss-positive Matses individual clusters with crAssphage strains from European individuals from the MetaHIT cohort, suggesting that it belongs to a widespread crAssphage clade.

To assess whether the lack of biogeographic structure in the multi-gene phylogeny was due to lack of sufficient information for phylogenetic analysis, we identified a subset of 118 samples fulfilling the following criteria: 1) according to the *de novo* assembly results, none of crAssphage proteins were matched by more than one sample ORF, and 2) according to the reference-based mapping, more than 70% of the crAssphage genome was covered at least 10-fold. Furthermore, we determined that these strains showed a range of 2 – 280 heterozygous sites (i.e. a maximum of 0.3% of sites across the genome were heterozygous). Taken altogether, we considered these samples to contain only one dominant crAssphage strain. The reference crAssphage genome was also included. A multi-genome alignment comprising all 97,065 nucleotide sites was built and used to generate an ML tree (Fig 1). This phylogeny supports the findings of our multi-gene analysis. CrAssphage strains from the same individual cluster together, with high bootstrap support. Strains from individuals from the same study dataset do not all necessarily cluster together; many such phylogenetically dissimilar strains are found in individuals within a study population. At the level of broadly defined geographic regions, some biogeographic structure is observed. However, even at phylogenomic resolution, due to low bootstrap values, the relative positions of these clusters within the crAssphage phylogeny cannot be well-defined.

**Fig 1.**
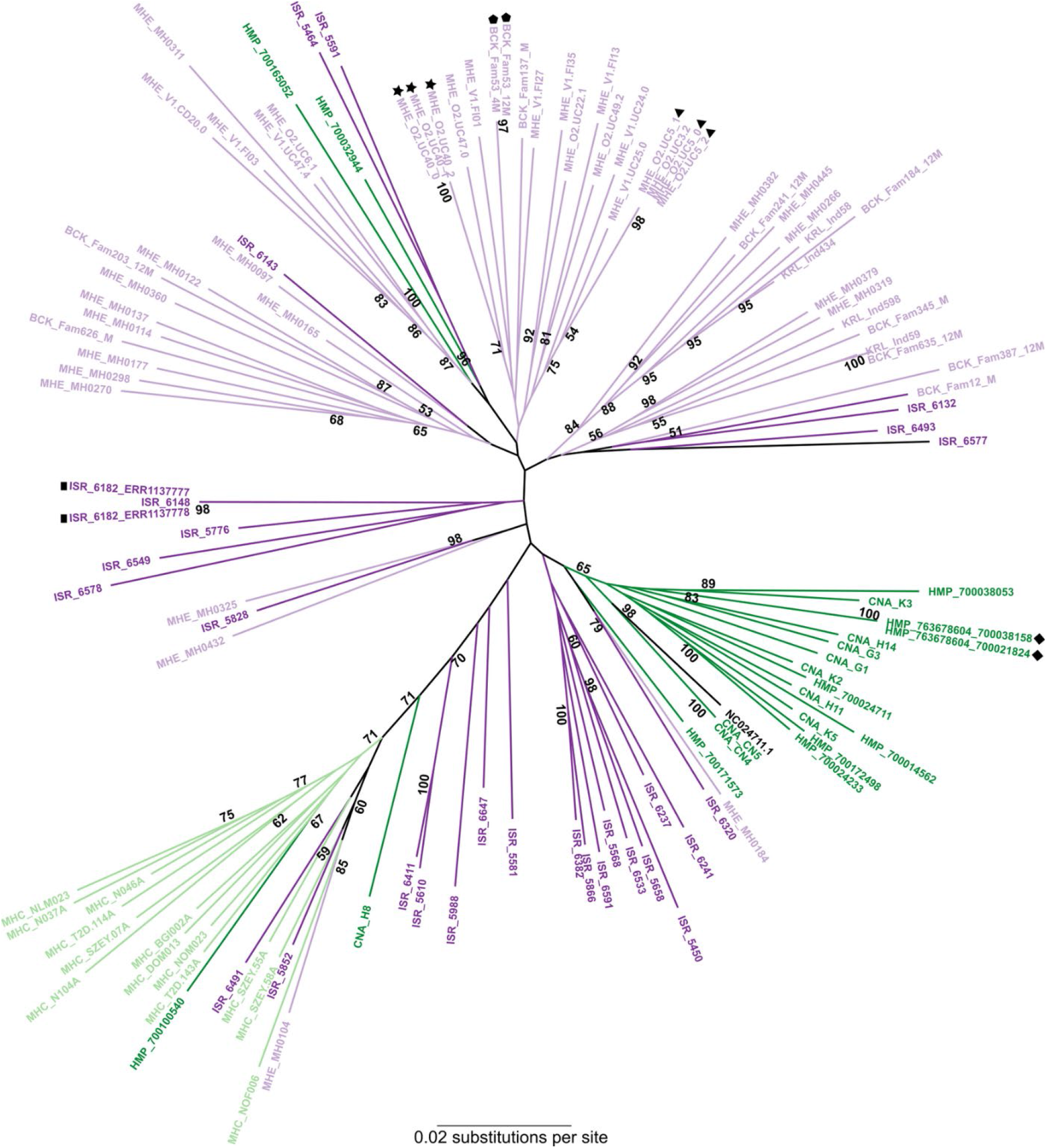
Phylogenomic analysis of crAssphage strains. The Maximum Likelihood tree was based on a multi-genome alignment comprising 97,065 sites. The tree was built using the Generalized Time-Reversible model with gamma-distributed rate variation and proportion of invariant sites. Bootstrap support was estimated from 100 replicates; only values greater than 50% are shown. Strains are color-coded according to the geographic location: dark green – The Americas, light green – Asia, light violet – Europe, and dark purple – Middle East (Israel). Symbols are used to denote strains from the same individual.

### CrAssphage population expansion

We hypothesized that the lack of strong biogeographic clustering could be explained by a recent expansion event in crAssphage. To asses this, we used a Bayesian Evolutionary Analysis Sampling Trees (BEAST) approach [47]. The analysis was performed using the multi-gene alignment described previously, and a fixed mean substitution rate, since these crAssphage strains are mostly contemporaneous. Because a substitution rate for crAssphage or other *Podoviridae* bacteriophages has not been estimated as of yet, we used a rate of 1.9 x 10^-4^ substitutions per site per year as estimated for bacteriophages in the family *Siphoviridae* [48], the members of which also infect bacteria and archaea. Our Bayesian Skyline plot (BSP) (S3 Fig) shows that crAssphage strains underwent an expansion event likely within the past 200 years. However, we stress the uncertainty of this estimate given the limitations of our Bayesian dating approach – namely, the lack of appropriate external calibration points and a substitution rate specific to crAssphage. Given the recent successful propagation of crAssphage in culture [44], the determination of this substitution rate has become a possibility. Further investigation using a crAssphage-specific substitution rate will provide a more robust estimate of the timing of the population expansion.

### CrAssphage acquisition and persistence

Acquisition and persistence of crAssphage strains in the human gut was evaluated using data from studies focusing on 1) mother-infant pairs, 2) twin-pairs, and 3) longitudinal sampling of individuals. We assessed the potential to identify vertical transmission of crAssphage using the BCK dataset [22], which comprises samples from healthy Swedish mothers (N=100) and their infants (N=100) at birth, 4-months, and 12-months of age. CrAssphage prevalence among the healthy mothers was 23%. None of the samples from newborns were considered crAss-positive. Most of the samples from newborn infants showed no reads mapping to crAssphage and no crAssphage proteins recovered from the *de novo* metagenomic assembly approach (S2 Table), with the exception of the infants from Family 549 and Family 263, who showed presence of three and five crAssphage proteins, respectively, at birth. The mothers from both families were also crAss-positive. The infant from Family 263 was determined to be crAss-positive at the four- and 12-month stages, whereas the infant from Family 549 was never or Ass-positive, despite showing a steady increase in the number of crAssphage proteins recovered. At the four-month stage, a total of three infants were crAss-positive, all of whom remained crAss-positive at the 12-month stage as well. A total of 16 infants were crAss-positive at the 12-month stage, suggesting that by the end of the first year of life, crAssphage prevalence among infants was similar to that in mothers (chi-square=1.5608; p-value=0.2115, α=0.05). Interestingly, in nine out of 16 mother-infant pairs, the 12-month old infants were crAss-positive whereas mothers were completely crAss-negative (no crAssphage proteins were recovered from the samples from the mothers). This supports the hypothesis that crAssphage may be acquired by means other than vertical transmission, as recently reported by [21].

To study concordance of crAssphage prevalence among twin-pairs, we screened the XIE dataset comprising samples from twin-pairs from the UK (N=124 pairs). We found 15 cases wherein both twins were crAss-positive and 57 cases wherein both twins were crAss-negative. There were 51 twin-pairs with discordant crAss-positive status. CrAssphage strains from only two crAss-positive twin-pairs could be included in our phylogenetic analyses: in one case, the strains from twin-pair P98 clustered together, whereas those from twin-pair P122 did not (Fig 2).

**Fig 2.**
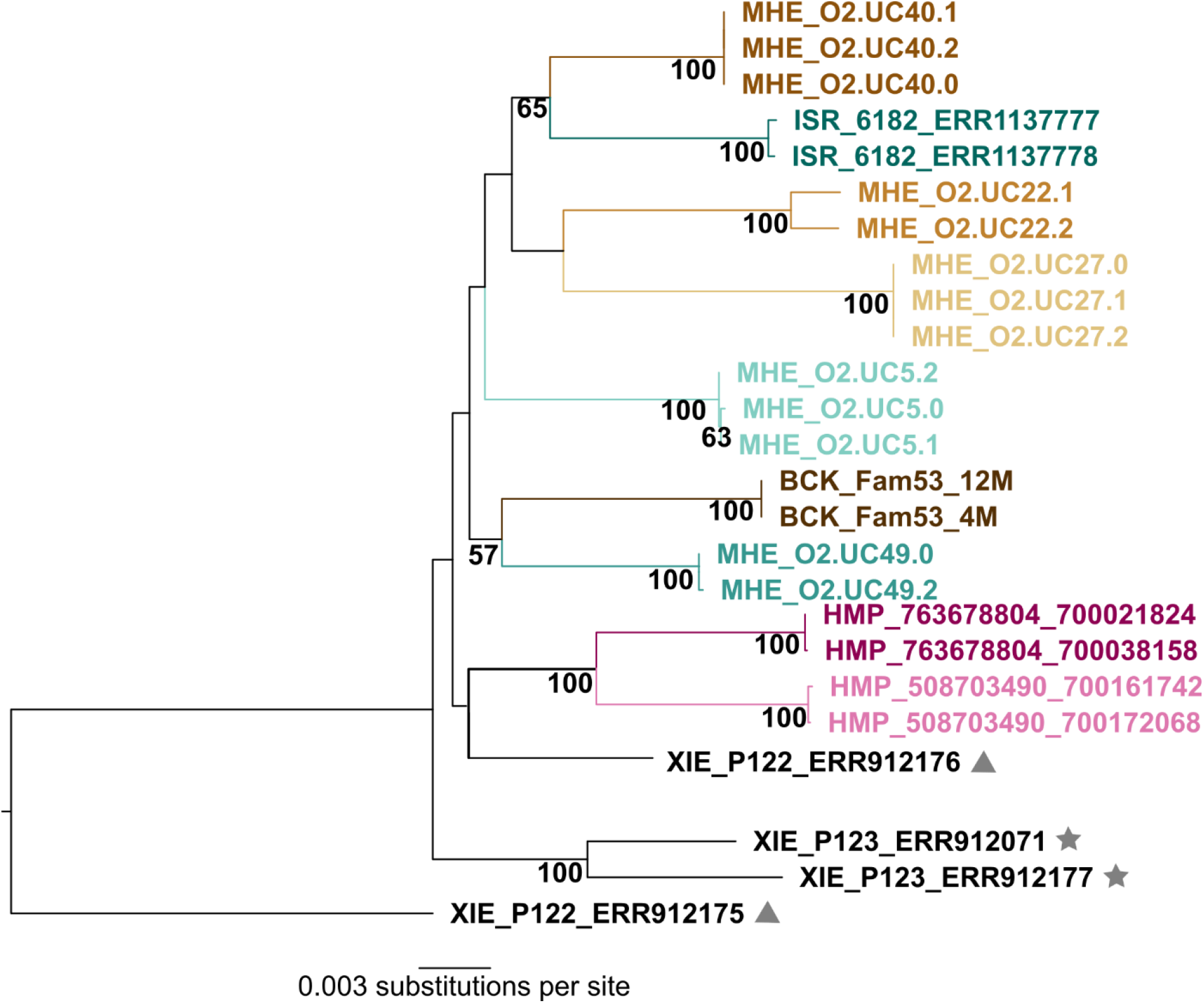
Relationships of crAssphage strains recovered from the same individual and twin-pairs. The Maximum Likelihood tree was based on the multi-gene alignment comprising 12,642 nucleotide sites. Sites with missing data were eliminated. The tree was built using the Generalized Time-Reversible model with gamma-distributed rate variation and proportion of invariant sites. Bootstrap support values estimated from 100 replicates are given; only values greater than 50% are shown. CrAssphage strains recovered from multiple samples from the same individual are color-coded accordingly (MHE, ISR, HMP, and BCK datasets). Symbols are used to denote crAssphage strains recovered from the same twin-pair (XIE dataset).

Additionally, we looked for signatures of crAssphage strain continuity among individuals using the BCK, HMP, and MHE datasets. As seen in Fig 2, crAssphage strains recovered at different time-points from the same individual cluster together, as also reported by [20]. In certain individuals (for example, MHE_O2.UC40), crAssphage strains from different time-points are a 100% identical, suggesting persistence of a single crAssphage strain. In others (for example, MHE_O2.UC22), the strains are closely related but not identical, suggesting that at different time-points, different crAssphage strains might be dominant in an individual.

## Conclusions

The geographic distribution of crAssphage is global [13, 20], but as observed here, the prevalence of crAssphage is lower within samples from more traditional, hunter-gatherer populations such as the Hadza from Tanzania and Matses from Peru. The overall picture from the data presented here is that crAssphage prevalence is associated with an industrialized lifestyle/diet, but with no associations to health, age, sex, or body-size variables. CrAssphage strains from the same individual tend to cluster together phylogenetically. Overall, crAssphage shows limited biogeographic clustering as seen in cases of a recent population expansion event. We estimate that this expansion occurred approximately within the past 200 years; however, the mechanism behind this expansion remains uncertain.

## Methods

### Data acquisition and processing

Gut metagenomic data for a total of 3,341 samples from were downloaded from the Sequence Read Archive or European Nucleotide Archive (Table 1). Shotgun data were processed using AdapterRemoval v2 [49] to remove reads with ambiguous bases (‘N”), trim at low-quality bases (Q<30), and merge overlapping read-pairs. Processed reads longer than 30 base pairs (bp) were retained for downstream analysis. These “analysis-ready” reads were then screened for the presence of crAssphage.

### Reference-based mapping

The analysis-ready reads were mapped to the reference crAssphage genome (NC024711.1) using Bowtie2 [37] with the “no-unal” option to discard unmapped reads. The resulting SAM files were processed using SAMTools v1.3 [50], converted into BAM files, quality-filtered at Phred threshold 37, and duplicate reads were removed using rmdup. SAMTools mpileup and VarScan v2.4.3 [51] were used to generate a VCF file with the following parameters: minimum coverage: 10, minimum coverage of variant allele: 3; minimum average quality: 30, minimum variant allele frequency: 0.2, minimum frequency for homozygotes: 0.9, p-value: 1, and strand filter: 0. This VCF file comprised both variant and invariant sites present in the reference crAssphage genome, resulting in a total of 97,065 sites. A custom perl script was used to generate a FASTA file containing the complete genome of the crAssphage strain from the VCF file. The number heterozygous sites was used to assess presence of one or multiple crAssphage strains in the sample.

### *De novo* metagenomic assembly

Analysis-ready reads were assembled into contigs using MEGAHIT [38]. Depth of coverage was calculated by mapping analysis-ready reads to assembled contigs using Bowtie2 [37], followed by processing of resulting alignment files using SAMTools [50] and custom R scripts. Open reading frame (ORF) prediction was carried out using Prodigal [52]. A custom BLAST database was created using the collection of 90 proteins previously predicted from the crAssphage reference genome (NC024711.1) [11]. The predicted ORFs (amino acid) from each shotgun metagenome were queried against this custom database using BLASTP [39], and matches were identified using the following criteria: 1) query coverage (length of alignment / query length) ≥ 95%, 2) percent identity ≥ 95%, and 3) E value < 1e-5. A sample was considered crAss-positive if matches were recovered for at least 33 reference crAssphage proteins.

### Association of crAssphage prevalence with metadata variables

Individuals were grouped on the basis of 1) health status, 2) age: < 18 years, 18 – 40 years, 41 – 65 years, and > 65 years, 3) sex: male and female, and 4) BMI: underweight (BMI < 18.5), normal (BMI 18.5 – 24.99), and overweight (BMI ≥ 25), according to the World Health Organization recommendations. Associations between the prevalence of crAssphage and health status, age, sex, and BMI categories were assessed using the Chi square test in R [53].

### Gut bacterial taxonomic profiles

For the MAT dataset [31], analysis-ready reads for each individual were mapped to the Greengenes database of 16S rRNA gene sequences [54] using Bowtie2 [37]. Unmapped reads were removed using the --no-unal option. The resulting SAM files were converted to BAM files, sorted, and duplicates were removed using SAMTools v1.3 [50]. The sequences of the reads mapping to the Greengenes database were obtained from the BAM files. These sequences were clustered into Operational Taxonomic Units (OTUs) using vsearch [55], employing the cluster_fast algorithm and comparing to the GreenGenes database. Other parameters used included a minimum sequence length of 70 bp and 97% similarity for clustering. The resulting OTU table was rarefied to a depth of 10,000 sequences per sample and singleton OTUs were removed using QIIME [56]. Taxonomic summaries were generated at the phylum- and genus-levels. Relative abundances of genus *Bacteroides* in individuals were determined and plotted using R [53].

### Multi-gene phylogenetic analysis

For each assembled metagenome, the number of unique ORFs matching each crAssphage reference protein was calculated from the BLASTP results (S2 Table) and used to infer the number of concurrent crAssphage strains carried in the sample. Only samples identified as containing one unique ORF matching each crAssphage reference protein were selected for phylogenetic analysis. Depth of coverage information was used to verify that identified ORFs were representative of the same viral genome.

A previously published study demonstrated the use of putative capsid proteins (genes 75, 76, 77, 78, and 79) to document diversity among crAss-like phages [11]. Multi-gene phylogenetic analyses were conducted using these capsid protein-encoding genes, as well as five other genes found in members of the crAssphage family. These included genes encoding hypothetical proteins (gene 20 and gene 23), single-stranded DNA-binding protein (gene 21), DNA-G family primase (gene 22), and PD-(D/E)XK family nuclease (gene 85). A subset of 232 samples and the reference crAssphage genome was included in this analysis. The gene sequences were aligned separately using MUSCLE v.3.8.31 with default parameters [57] and concatenated together. Sites with missing data and gaps were completely removed. A Maximum Likelihood (ML) tree was built using RAXML v8.2.4 [58], using the Generalized Time-Reversible model with gamma-distributed rate variation and proportion of invariable sites (GTR+G+I). Bootstrap support was estimated from 100 replicates.

### Genome-based phylogenetic analysis

A subset of 118 samples determined to carry only one crAssphage strain was selected. A whole-genome alignment of all strains was used as input for RAXML [58], and a ML tree was generated using the GTR+G+I model and 100 bootstrap replicates.

### Timing of crAssphage population expansion

To estimate the timing of a potential population expansion among crAssphage sequences, the multi-gene alignment was used as input for BEAST v1.8.4 [47]. Since substitution rates for crAssphage or crAss-like phages remain undetermined, we used a substitution rate of 1.9 x 10^-4^ substitutions per site per year as estimated for bacteriophages belonging to the *Siphoviridae* family [48]. We used a GTR+G+I model of nucleotide substitution, an uncorrelated lognormal clock model with uniform rate across branches, and a Bayesian Coalescent Skyline plot tree prior. One Markov Chain Monte Carlo (MCMC) run was carried out with 100,000,000 iterations, sampling every 10,000 steps. The first 10,000,000 iterations were discarded as burn-in. Tracer [59] was used to visualize the results of the MCMC run and generate a Bayesian Skyline Plot (BSP).

## Supporting information

**S1 Table.** S1 Table contains a list of all 3,341 samples included in this study, associated metadata, and results of the crAssphage screening using reference-based mapping and *de novo* metagenomic assembly.

**S2 Table.** S2 Table shows the hits for all crAssphage reference proteins across the samples.

**S1 Fig. Relative abundance of genus *Bacteroides* among Matses individuals.** Boxplot denoting percentage relative abundance of *Bacteroides* sp. among Matses individuals. Values corresponding to crAss-positive individuals are denoted in red.

**S2 Fig. Multi-gene phylogenetic analysis of 233 crAssphage strains.** The Maximum Likelihood tree was based on the multi-gene alignment comprising 12,642 nucleotide sites. Sites with missing data were eliminated. The tree was built using the Generalized Time-Reversible model with gamma-distributed rate variation and proportion of invariant sites. Bootstrap support values estimated from 100 replicates are given; only values greater than 50% are shown. Strains are color-coded according to the geographic location: dark green – The Americas, light green – Asia, light violet – Europe, and dark purple – Middle East (Israel). The differentially-colored symbols next to the taxa names are used to denote strains from the same individual.

**S3 Fig. Bayesian Skyline Plot of crAssphage strains.** The X-axis denotes time in years before present (YBP) and Y-axis denotes estimated effective population size. The blue shaded region denotes the 95% Highest Posterior Density interval.

