## Supplementary figures and images for "Biogeographic study of human gut-associated crAssphage suggests impacts from industrialization and recent expansion"

### F1 Fig

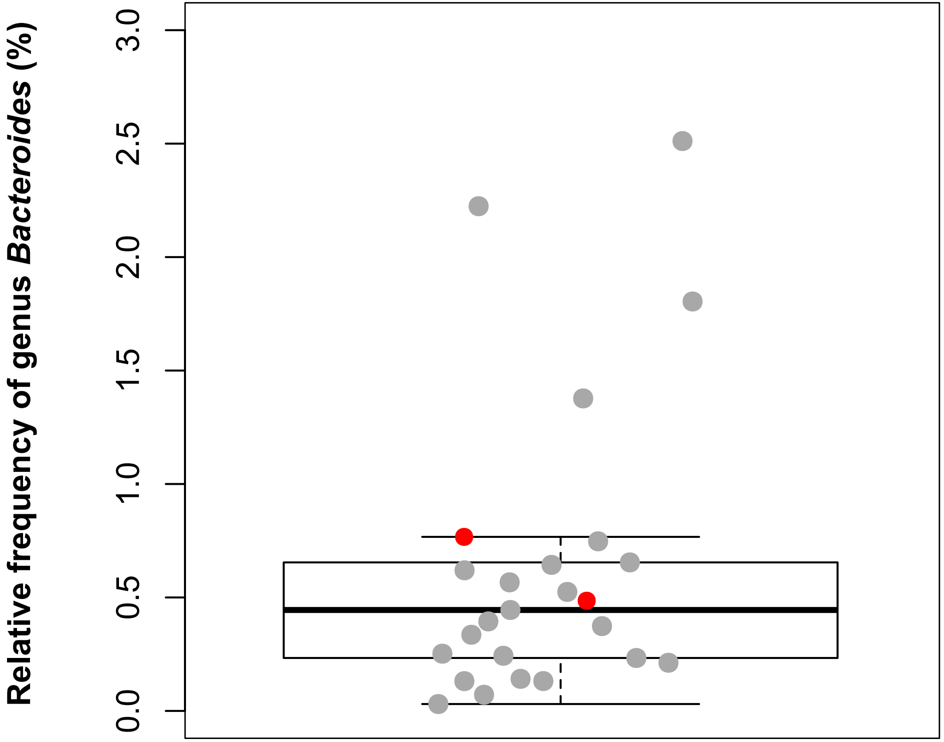

### S2 Fig

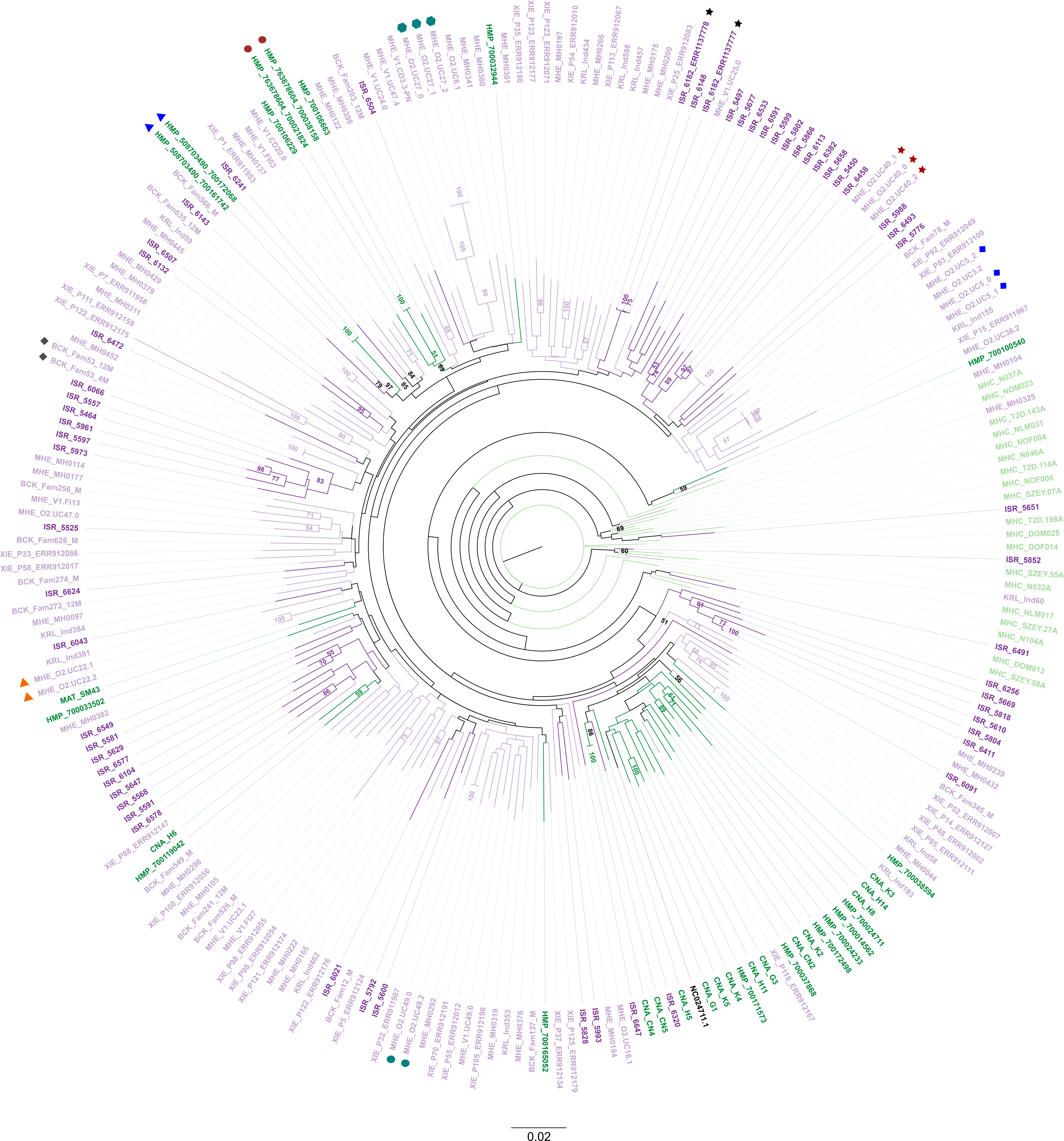

### S3 Fig

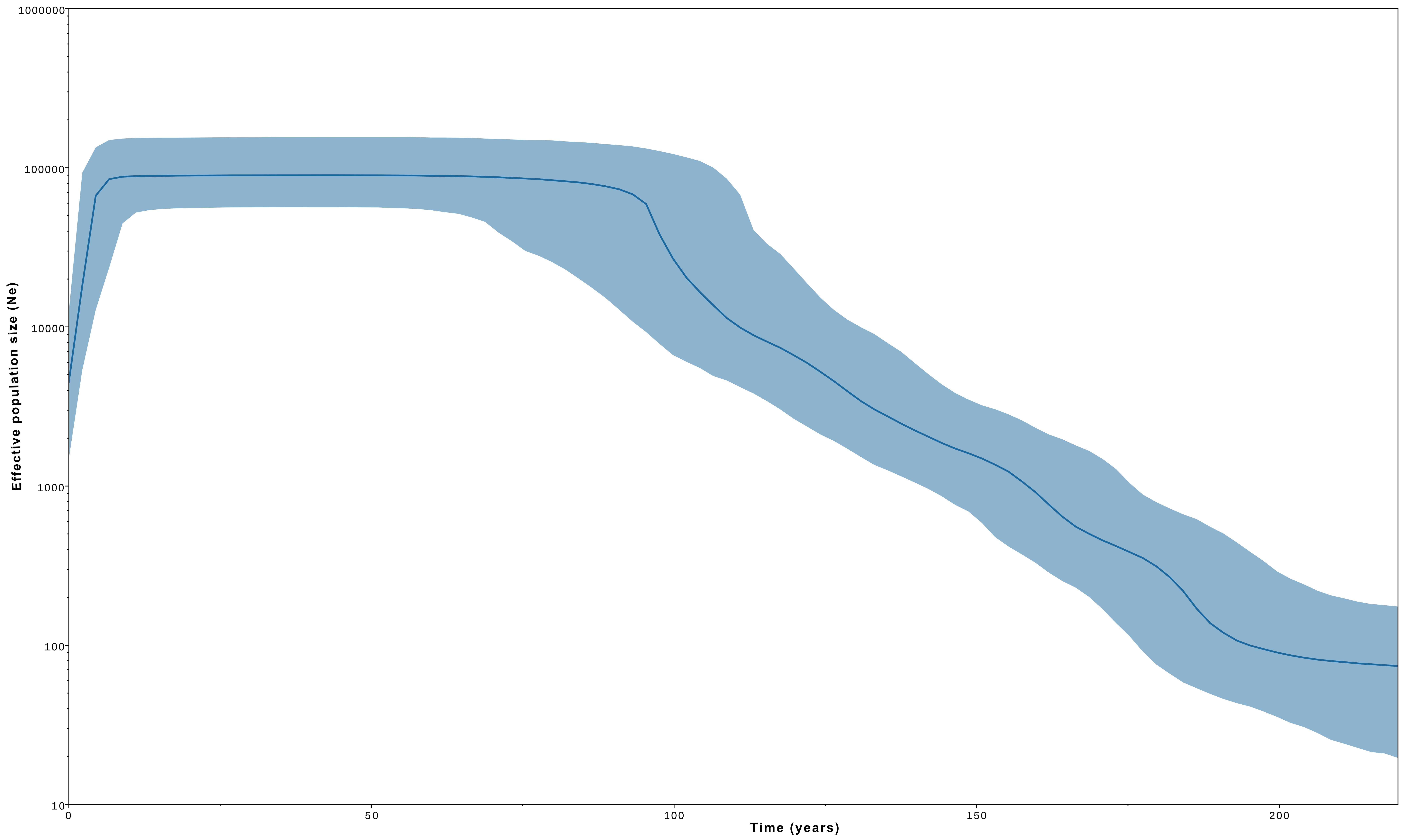
